# A Web-scraped Skin Image Database of Monkeypox, Chickenpox, Smallpox, Cowpox, and Measles

**DOI:** 10.1101/2022.08.01.502199

**Authors:** Towhidul Islam, Mohammad Arafat Hussain, Forhad Uddin Hasan Chowdhury, B. M. Riazul Islam

**Affiliations:** Department of Computer Science and Engineering, Northern University Bangladesh, Dhaka, 1215, Bangladesh; Department of Medicine, Dhaka Medical College Hospital, Dhaka, 1000, Bangladesh; Health Information Unit, Directorate General of Health Services, Dhaka, 1212, Bangladesh

**Keywords:** Monkeypox, Chickenpox, Smallpox, Cowpox, Measles, machine learning, deep learning, image data, skin lesions, CNN

## Abstract

Monkeypox has emerged as a fast-spreading disease around the world and an outbreak has been reported in 75 countries so far. Although the clinical attributes of Monkeypox are similar to those of Smallpox, skin lesions and rashes caused by Monkeypox often resemble those of other types of pox, for example, chickenpox and cowpox. This scenario makes an early diagnosis of Monkeypox challenging for the healthcare professional just by observing the visual appearance of lesions and rashes. The rarity of Monkeypox before the current outbreak further created a knowledge gap among healthcare professionals around the world. To tackle this challenging situation, scientists are taking motivation from the success of supervised machine learning in COVID-19 detection. However, the lack of Monkeypox skin image data is making the bottleneck of using machine learning in Monkeypox detection from patient skin images. Therefore, in this project, we introduce the Monkeypox Skin Image Dataset 2022, the largest of its kind so far. We used web-scraping to collect Monkeypox, Chickenpox, Smallpox, Cowpox, and Measles infected skin as well as healthy skin images to build a comprehensive image database and make it publicly available. We believe that our database will facilitate the development of baseline machine learning algorithms for early detection of Monkeypox in clinical settings. Our dataset is available at the following Kaggle link: https://www.kaggle.com/datasets/arafathussain/monkeypox-skin-image-dataset-2022.

## 1 Introduction

While the world is gradually recovering from the havoc caused by Coronavirus disease (COVID-19), another infectious disease, known as Monkeypox, has been spreading around the world at a rapid pace. To date, an outbreak of Monkeypox has been reported in 75 countries.^1^ Human infection of the Monkeypox virus was first reported in the Democratic Republic of Congo (formerly Zaire) in 1970 (Thornhill et al, 2022), which was transmitted to humans from animals. The monkeypox virus belongs to the genus *Orthopoxvirus* of the family *Poxviridae* (Shchelkunov et al, 2002), which shows symptoms similar to those of Smallpox (Thornhill et al, 2022). Smallpox was eradicated in 1970, leading to the cessation of Smallpox vaccination. Since then, Monkeypox has been considered the most important *Orthopoxvirus* for human health. Monkeypox used to be primarily reported in the African continent, however, it has been widely spreading in urban areas in different locations of the world (Thornhill et al, 2022). The current outbreak of Monkeypox in humans on a global scale is believed to be due to changes in Monkeypox’s biological attributes, changes in human behavior, or both.^2^

The clinical attributes of Monkeypox are similar to that of Smallpox.^3^. Additionally, Monkeypox skin lesions and rashes often resemble those of other types of pox, for example, chickenpox and cowpox, making an early diagnosis of Monkeypox challenging for the healthcare professional. The rarity of Monkeypox before the current outbreak (Sklenovska and Van Ranst, 2018) also created a knowledge gap among healthcare professionals around the world. Polymerase chain reaction (PCR) is typically considered the most accurate tool for the Monkeypox detection (Erez et al, 2019), however, healthcare professionals are accustomed to diagnosing pox infections by visual observation of skin rash and lesion. Despite a low mortality rate from Monkeypox infection (i.e., 1%-10%) (Gong et al, 2022), early detection of Monkeypox can facilitate patient isolation, as well as contact tracing for effective containment of community spread of Monkeypox.

Different machine learning (ML), especially deep learning (DL), approaches have been used in different medical image analysis tasks (e.g., organ localization (Hussain et al, 2021a, 2017), organ abnormality detection (Hussain et al, 2016, 2017), gene mutation detection (Hussain et al, 2018), cancer grading (Hussain et al, 2021b, 2019a) and staging (Hussain et al, 2019b)), and recently played a significant role in COVID-19 detection and severity analysis from images of different medical imaging modalities (e.g., computed tomography (CT), chest X-ray, and chest ultrasound) (Sun et al, 2022; Akbarimajd et al, 2022; Momeny et al, 2021). This success motivates the use of ML or DL approaches to detect Monkeypox from the skin images of patients. However, supervised or semi-supervised learning approaches are data-driven and require a large number of data to well-train an ML or DL model. Unfortunately, there is no publicly available healthcare facility or infectious disease control authority-disclosed database of Monkeypox skin lesions or rash. In such a situation, web-scraping (i.e., extracting data from websites) (Dogucu and Çetinkaya-Rundel, 2021) to collect Monkeypox skin lesion images may be the only alternative to facilitate the development of ML- and DL-based Monkeypox infection detection algorithms.

In this project, we used web-scraping to collect Monkeypox, Chickenpox, Smallpox, Cowpox, and Measles infected skin as well as healthy skin images to build a comprehensive image database and made it publicly available. To date, to our knowledge, there are two other Monkeypox databases based on web-scraping that are currently available (Ahsan et al, 2022; Ali et al, 2022). However, the number of data and the number of pox classes are limited in those databases. Our database is unique in the following way:

1. Our database contains skin lesion/rash images of five different diseases, that is, Monkeypox, Chickenpox, Smallpox, Cowpox, and Measles, as well as contains healthy skin images.
2. This database contains more web-scrapped pox and measles image data, before augmentation, compared to other similar databases.
3. We improved the privacy of patients in images by blocking exposed eyes and private parts with black boxes.
4. We used various augmentation techniques to increase the number of data by 49-times.
5. We acknowledged the sources of all images in our database and presented the source/credit list as supplementary material.

## 2 Methodology

In this section, we briefly describe our approach to data collection, expert screening, and data augmentation. We also show the pipeline of our database development in Fig. 1.

**Fig. 1:**
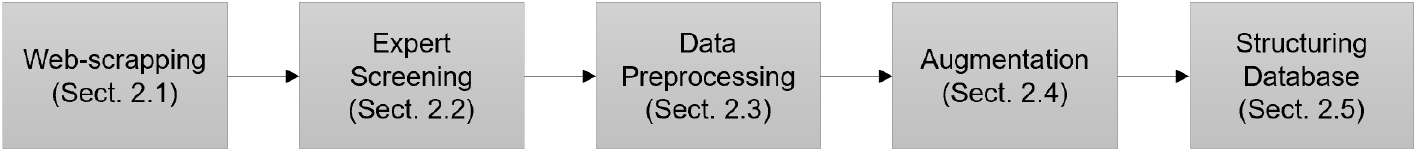
The pipeline of our database development. Each block mentions the corresponding section number in this paper, where the details are presented.

### 2.1 Web-scraping for Data Collection

As we mentioned in section 1 that a publicly available healthcare facility or infectious disease control authority-disclosed database of Monkeypox skin lesions or rashes is yet to come, we used web-scraping to collect Monkeypox, Chickenpox, Smallpox, Cowpox, and Measles infected skin as well as healthy skin images. We use the Google search engine to search for different types of pox-infected skin images, Measles-infected skin images, and healthy skin images from various sources such as websites, news portals, blogs, and image portals. We modified our search to collect images that fall under “Creative Commons licenses.” However, for several pox classes, we hardly find images under “Creative Commons licenses,” thus collected images that fall under “Commercial & other licenses.” Therefore, we include a list as supplementary material that includes the uniform resource locator (URL) of the source, access date, and photo credit (if any) for all our collected images. In Fig. 2, we show some example images from our database.

**Fig. 2:**
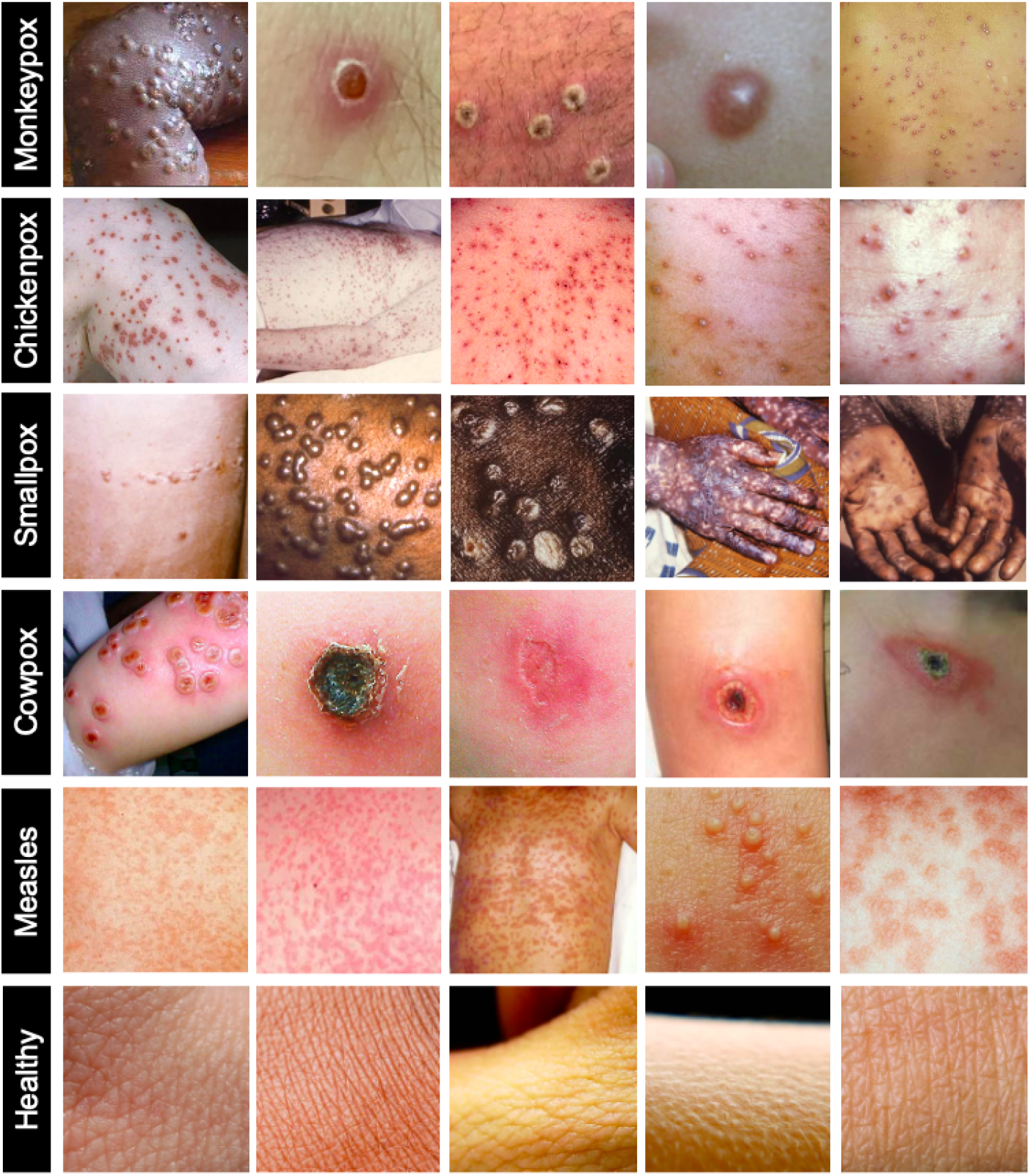
Example skin images of Monkeypox, Chickenpox, Smallpox, Cowpox, Measles, and healthy cases (first to sixth rows, respectively) from our database.

### 2.2 Expert Screening of Data

After collecting image data, two expert physicians, who specialized in infectious diseases, sequentially screened all the images to validate the supposed infection. In Fig. 3, we show a pie chart of the percentage of original web-scrapped image data per class in our database, after the expert screening.

**Fig. 3:**
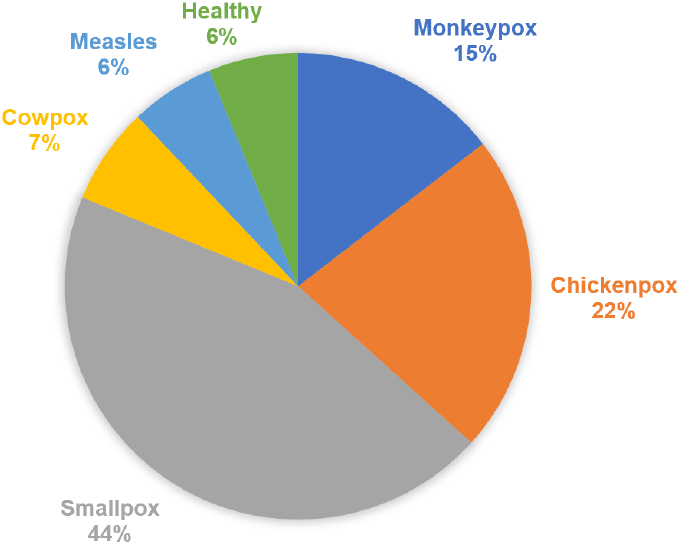
A pie chart showing the percentage of original web-scrapped image data per class in our database.

### 2.3 Data Preprocessing

In this step, we manually cropped images to remove unwanted peripheral background regions. Furthermore, to make patients nonidentifiable from their corresponding images, we blocked the eye region with black boxes. We also did the same to hide revealed private parts as much as possible, without blocking the skin lesions/rashes. Since typical DL algorithms expect a square-shaped input image in terms of pixel counts (often 224×224×3 pixels), we added extra blank pixels in the periphery of many images to avoid excessive stretching of the actual skin lesions during image resizing. After that, we cropped and resized all the images to 224*×*224*×*3 pixels (3-channel RGB format).

### 2.4 Augmentation

To increase the number of images and introduce variability in the data, we performed the following 19 augmentation operations on the web-scrapped image data using Python Imaging Library (PIL) version 9.2.0, and scikit-image library version 0.19.3:

1. Brightness modification with a randomly generated factor (range [0.5, 2]).
2. Color modification with a randomly generated factor (range [0.5, 1.5]).
3. Sharpness modification with a randomly generated factor (range [0.5, 2]).
4. Image translation along height and width with a randomly generated distance between -25 and 25 pixels.
5. Image shearing along height and width with randomly generated parameters.
6. Adding Gaussian noise of zero mean and randomly generated variance (range [0.005, 0.02]) to images.
7. Adding zero-mean speckle noise and randomly generated variance (range [0.005, 0.02]) to images.
8. Adding salt noise to randomly generated number of image pixels (range [2%, 8%]).
9. Adding pepper noise to the randomly generated number of image pixels (range [2%, 8%]).
10. Adding salt & pepper noise to randomly generated number of image pixels (range [2%, 8%]).
11. Modifying the values of the image pixels based on the local variance.
12. Blurring an image with a Gaussian kernel with a randomly generated radius (range [1, 3] pixels).
13. Contrast modification with a randomly generated factor (range [1, 1.5]).
14. Rotating all images by 90^*°*^.
15. Rotating images at random angle (range [-45^*°*^, 45^*°*^]).
16. Zooming in an image by about 9%.
17. Zooming in an image by about 18%.
18. Flipping images along the height.
19. Flipping images along the width.

In Table 1, we show the number of original and augmented images per class in our database. We also show a flow diagram of our augmentation pipeline in Fig. 4. We use these augmentation operations to increase the data by 49*×*.

**Table 1:** Distribution of image classes in the Monkeypox Skin Image Dataset.

| Class label | No. of Original Images | No. of Augmented Images |
| --- | --- | --- |
| Monkeypox | 117 | 5,733 |
| Chickenpox | 178 | 8,722 |
| Smallpox | 358 | 17,542 |
| Cowpox | 54 | 2,646 |
| Measles | 47 | 2,303 |
| Healthy | 50 | 2,450 |
| **Total** | **804** | **39,396** |

**Fig. 4:**
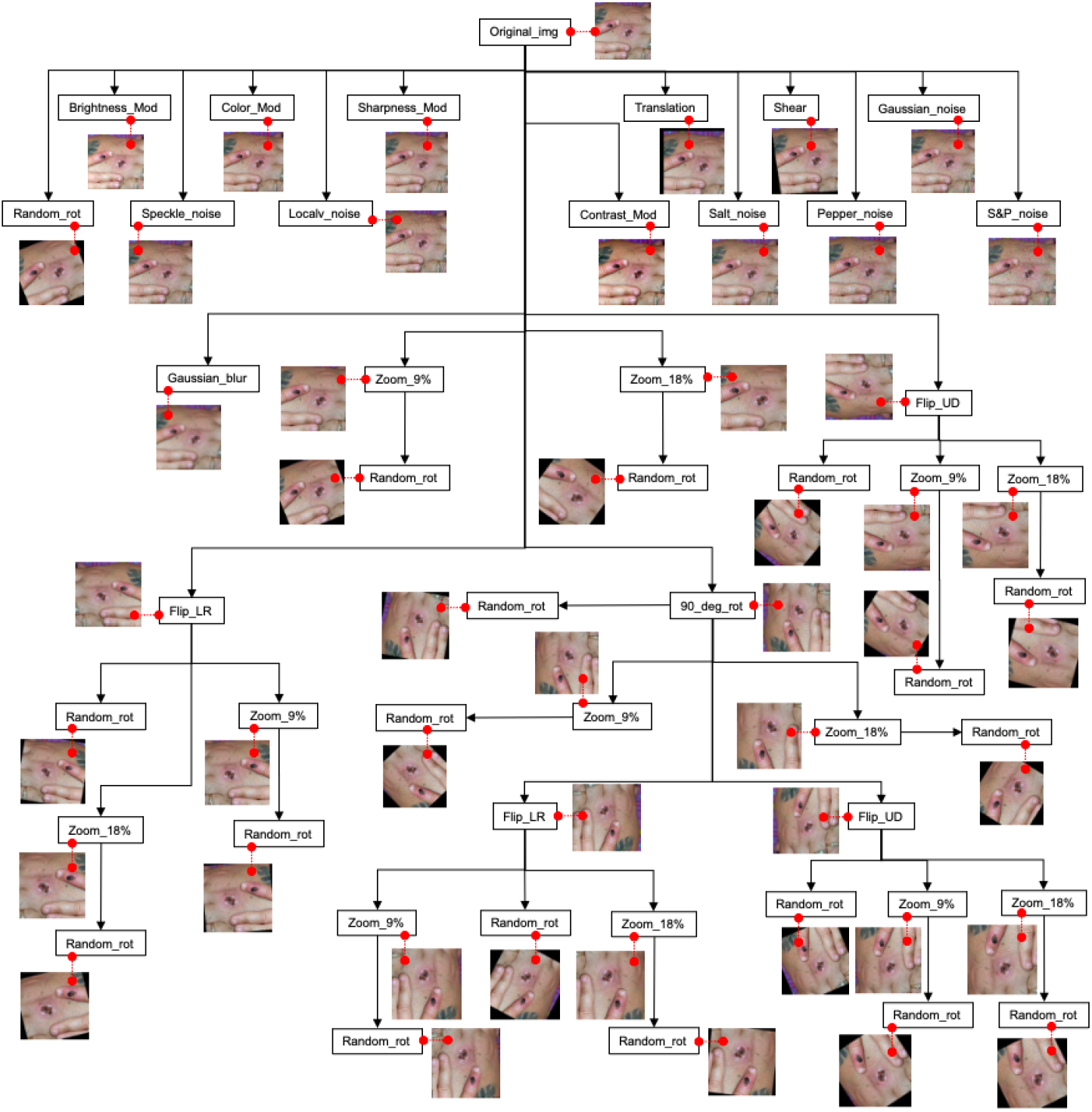
Flow diagram of our augmentation pipeline. Using this pipeline of operations, we increased the data by 49×. We also show representative augmented images attached to each box. Acronyms- Original_img: original image, Brightness_Mod: brightness modification, Color_Mod: color modification, Sharpness_Mod: sharpness modification, Gaussian_noise: Gaussian noise addition, Random_rot: random rotation, Speckle_noise: Speckle noise addition, Localv_noise: local variance noise addition, Contrast_Mod: contrast modification, Salt_noise: salt noise addition, Pepper_noise: pepper noise addition, S&P_noise: salt & pepper noise addition, Gaussian_blur: Gaussian blurring, Zoom_9%: 9% zooming in, Zoom_18%: 18% zooming in, Flip_LR: flipping along the width, Flip_UD: flipping along the height, and 90_deg_rot: rotation of image by 90^*°*^.

### 2.5 Naming Convention and Database Structure

We named the pre-processed original images as “xx_yyyy.jpg,” where “xx” is either of “mo,” “ch,” “sm,” “co,” “me,” or “he,” representing Monkeypox, Chickenpox, Smallpox, Cowpox, Measles, or healthy, respectively, and “yyyy” denotes the serial of an image belonging to “xx” class. After augmentation, images in the pool are named “aug_xx_yyyy_zzzz.jpg,” where “zzzz” represents the serial of the augmented image of original image “yyyy” belonging to “xx” class. Note that “aug_xx_yyyy_0001.jpg” is always the preprocessed but resized original image. So, during validation and testing/inference, it will be sufficient to call “aug_xx_yyyy_0001.jpg” for accuracy estimation. Furthermore, during data partitioning for training, validation, and testing, or cross-validation, all “zzzz” of a particular “yyyy” should be in a specific fold (i.e., training, validation, or testing) to avoid cross-fold data splitting of the same patient “yyyy”.

We structured our database in a way that makes it easy to call in an ML or DL model training routine. In Fig. 5, we show the structure of our database in terms of directory map.

**Fig. 5:**
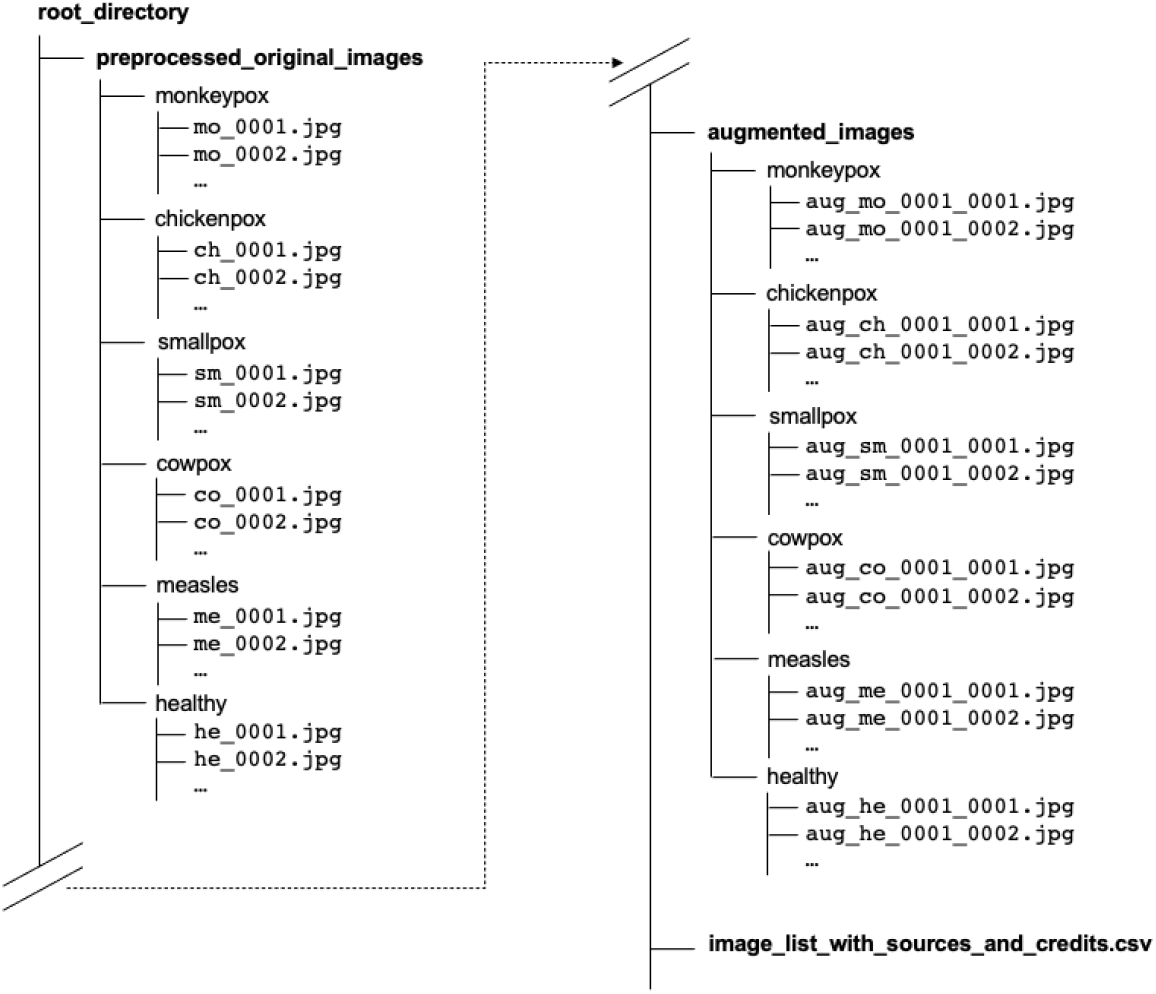
Directory map of our database.

## 3 Expected Outcomes

We expect that our proposed database will help to achieve the following objectives:

### Baseline ML and DL Algorithms

We expect that our database will facilitate the development of baseline ML and DL algorithms for early Monkeypox detection until a larger dataset from any national or international government or autonomous entity/authority comes out.

### Achieving Better Detection-ability

Unlike other similar databases, our database contains image data for more classes of pox, as well as Measles and healthy skin. Also, it contains more images per class than other databases. Therefore, we expect that training any ML or DL model on our dataset would achieve a better ability to classify different types of skin lesions/rashes.

### Avoiding Model Over-fitting

Since our database has multi-class images, which were again multiplied 49× by augmentation, we expect that any ML or DL model would be better generalized (i.e., not over-fitted on a specific disease class) if trained on our dataset.

### Aid in Clinical Environment

An ML or DL model, trained on our dataset, can help in the clinical diagnosis of Monkeypox. It may reduce the dependency on conventional microscopic image analysis and PCR-based Monkeypox detection. It may also reduce the close contact between healthcare professionals and patients. In fact, a patient can be diagnosed remotely by analyzing an image of the skin lesion of the patient taken with a smartphone.

## 4 Conclusion

In this project, we used web-scraping to build a comprehensive database of images of skin infected with Monkeypox, Chickenpox, Smallpox, Cowpox and Measles. Compared to other similar databases, ours has the largest number of actual images per class and the largest number of augmented images per class. We took the expert opinions of two physicians to validate the disease in an image. We also made a list to acknowledge the source of each image and made it available for public use and scrutiny. We believe that this database will facilitate the development of baseline ML and DL algorithms for early Monkeypox detection, enable ML or DL models to achieve better ability, as well as generalizability to classify different types of skin lesions/rashes, and overall help in the clinical diagnosis of Monkeypox. We also aim to keep this database updated with new data as soon as they appear on the web.

## Supporting information

List of data sources

## 5 Supplementary Material

The list of image sources for all classes can be found in this link: https://www.kaggle.com/datasets/arafathussain/monkeypox-skin-image-dataset-2022?select=image_list_with_sources_and_credits.xlsx

### Description of the Materials

- **File Name:** image_list_with_sources_and_credits.xlsx
- **Sheets:** There are six sheets in the spreadsheet. They are named as:
  1. chickenpox
  2. measles
  3. smallpox
  4. monkeypox
  5. cowpox
  6. healthy
- **Columns:** Each sheet has five columns as:
  1. Image Name (names of images as appear in the database)
  2. Source Link (the exact URL of the web page, from where the image has been adopted)
  3. Access Date (the date we accessed the image on a particular web page as mentioned in the second column)
  4. Source Name (the name of the hosting entity of the image)
  5. Photo Credit (if any; the person or organization, who captured the image)

## Footnotes

1 https://www.nytimes.com/2022/07/23/health/monkeypox-pandemic-who.html

2 https://www.cdc.gov/poxvirus/monkeypox/response/2022/world-map.html

3 https://www.who.int/news-room/fact-sheets/detail/monkeypox

